# DNAJB chaperones inhibit aggregation of destabilised proteins via a C-terminal region distinct from that used to prevent amyloid formation

**DOI:** 10.1101/2020.10.08.326280

**Authors:** Shannon McMahon, Steven Bergink, Harm H. Kampinga, Heath Ecroyd

## Abstract

Disturbances to protein homeostasis (proteostasis) can lead to protein aggregation and inclusion formation, processes associated with a variety of neurodegenerative disorders. DNAJBs are molecular chaperones previously identified as potent suppressors of disease-related protein aggregation. In this work, we over-expressed a destabilised isoform of firefly luciferase (R188Q/R261Q Fluc; Fluc^DM^) in cells to assess the capacity of DNAJBs to inhibit inclusion formation. Co-expression of all DNAJBs tested significantly inhibited the intracellular aggregation of Fluc^DM^. Moreover, we show that DNAJBs suppress aggregation by supporting the Hsp70-dependent degradation of Fluc^DM^ via the proteasome. The serine-rich stretch in DNAJB6 and DNAJB8, essential for preventing fibrillar aggregation, is not involved in the suppression of Fluc^DM^ inclusion formation. Conversely, deletion of the C-terminal TTK-LKS region in DNAJB8, a region not required to suppress polyQ aggregation, abolished its ability to inhibit inclusion formation by Fluc^DM^. Thus, our data suggest that DNAJB6 and DNAJB8 possess two distinct domains involved in the inhibition of protein aggregation, one responsible for binding to β-hairpins that form during amyloid formation and another that mediates the degradation of destabilised client proteins via the proteasome.

**Summary statement:** Specialised DNAJB molecular chaperones are potent suppressors of protein aggregation and interact with different types of client proteins via distinct C-terminal regions

## Introduction

Many age-related neurodegenerative diseases, including Alzheimer’s disease, Huntington’s disease and amyotrophic lateral sclerosis (ALS), are associated with the expression of aggregation-prone proteins or polypeptide fragments that oligomerise and deposit into inclusions. For example, the deposition of the tau protein and amyloid-β peptide into intracellular and extracellular deposits, respectively, are pathological hallmarks of Alzheimer’s disease (Bucciantini et al., 2002; Hardy and Selkoe, 2002; Haass and Selkoe, 2007; Iqbal et al., 2009). Proteins containing expanded polyglutamine (polyQ) repeats aggregate to from amyloid precursors and mature fibrils which have been linked to Huntington’s disease and various ataxias (Zoghbi and Orr, 2000; Chiti and Dobson, 2006). Although many diseases are characterised by the formation of amyloid fibrils, not all proteins aggregate through an ordered mechanism. For example, superoxide dismutase 1 (SOD1) mutations, which are causative of some familial forms of ALS, forms highly disordered and hydrophobic amorphous precipitates (Banci et al., 2007; Prudencio et al., 2009). In addition, recent studies have demonstrated that almost all cases of sporadic ALS and frontotemporal dementia share a common neuropathology of predominantly amorphous, intracellular deposits that contain TAR DNA-binding protein 43 (TDP-43) (Neumann et al., 2006; Adachi et al., 2009; Scotter et al., 2015). Furthermore, some proteins are known to adopt intermediate conformations before forming amyloid fibres (Dobson, 2003; Stathopulos et al., 2003). Finally, amorphous aggregation is associated with protein misfolding that occurs during conditions of cellular stress (Chiti and Dobson, 2006; Ecroyd and Carver, 2008). Thus, protein aggregation can lead to amyloid-fibril and/or amorphous forms, each with their own unique characteristics (Kampinga and Bergink, 2016): it is likely that different mechanisms or protein quality control machinery is required to maintain each of these forms of aggregation-prone proteins in a soluble state.

Cellular protein homeostasis (proteostasis) is maintained by an interconnected protein quality control (PQC) network that helps to maintain a stable and functional proteome. The overload or failure of these PQC systems to maintain the solubility and function of aggregation-prone proteins has been hypothesised to result in the inclusion formation and deposits associated with protein conformational diseases (Kampinga and Bergink, 2016). Molecular chaperones, in conjunction with various co-factors, including co-chaperones, are a key component of the PQC network. Molecular chaperones have a central role in ensuring the correct folding of nascent polypeptides, the maintenance of partially folded protein intermediates in a folding-competent state, the re-folding of damaged proteins and, where required, the shuttling of misfolded aggregation-prone proteins for degradation. The heat shock proteins (Hsps) are a family of evolutionarily conserved molecular chaperones. They were discovered as proteins whose expression and activity is dramatically upregulated in response to a variety of (mostly) proteotoxic forms of stress (Feder and Hofmann, 1999). Most of these stress-inducible Hsps bind to exposed hydrophobic regions of proteins and either assist in their refolding, or traffic irreversibly damaged proteins for proteolytic degradation via the proteasome or by autophagy (Powers et al., 2009; Morimoto, 2011).

Critical components of the chaperone network include the Hsp70 machinery. The activity of these Hsp70 chaperone machines is driven by interactions with DNAJ proteins, whereby DNAJs deliver misfolded substrates to the Hsp70 machinery and in turn, stimulate Hsp70 ATPase activity (Bukau et al., 2006). The human genome encodes 53 DNAJ proteins that can be divided into three separate subfamilies, the DNAJA, DNAJB and DNAJC proteins, based upon intrinsic structural features (Cheetham and Caplan, 1998; Kampinga and Craig, 2010). Of these, the DNAJB proteins are the most extensively studied due to them being previously identified as potent suppressors of aggregation associated with many disease-related proteins, including polyQ, amyloid-β and SOD1 proteins. For example, previous work has shown that DNAJB2a and the closely related members DNAJB6a/b and DNAJB8 potently suppress the aggregation of a polyQ-expanded protein in a cell culture model of disease (Howarth et al., 2007; Hageman et al., 2010; Gillis et al., 2013). Increasing the expression of DNAJB2a or DNAJB6b in a mouse model of Huntington’s disease delayed polyQ aggregation, alleviated symptoms and prolonged lifespan (Labbadia et al., 2012; Kakkar et al., 2016b). Moreover, DNAJB6b, both *in vitro* and in cells, prevents the nucleation of the amyloid-β peptides into mature fibrils (Månsson et al., 2014). More recently, mutant SOD1 aggregation was reported to be significantly suppressed by DNAJB1, DNAJB2b and DNAJB7, whilst other DNAJBs were found to have little or no effect (Serlidaki et al., 2020).

Structurally, DNAJB proteins contain a highly conserved N-terminal J-domain, an internal glycine/phenylalanine (G/F)-rich linker region and a C-terminal domain (Cheetham and Caplan, 1998). The J-domain contains the conserved histidine-proline-aspartate (HPD) motif for interaction with the Hsp70 machinery, whilst the C-terminal region is thought to bind substrates (Kampinga and Craig, 2010). Specialised members of the DNAJB family (DNAJB6 and DNAJB8) contain a serine/threonine (S/T)-rich motif in between the G/F-rich region and C-terminal domain. The hydroxyl groups within the side chains of this S/T-rich region participate in intramolecular hydrogen bonding with β-hairpin structures in amyloid-β and polyQ peptides to prevent their primary nucleation into mature (disease causing) toxic amyloid fibres (Hageman et al., 2010; Månsson et al., 2014; Kakkar et al., 2016b). The role of the G/F-rich region is currently not well understood; however, mutations in this region have been linked to reduced substrate binding capacity (Perales-Calvo et al., 2010) and have been implicated in inheritable forms of limb-girdle muscular dystrophy (Harms et al., 2012; Sarparanta et al., 2012). It has therefore been hypothesised that the G/F-rich region may also be directly involved in substrate binding as well as participate as a flexible linker region for inter-domain stabilisation.

To-date, the majority of cell-based studies conducted to reveal the interaction of DNAJBs with client proteins have investigated proteins whose aggregation is disease related (Hageman et al., 2010; Hageman et al., 2011; Månsson et al., 2014; Kakkar et al., 2016a; Serlidaki et al., 2020). In this work we have chosen to exploit a previously described mutant isoform of firefly luciferase (R188Q/R261Q Fluc; herein referred to as Fluc^DM^) (Gupta et al., 2011) to assess the generic capacity of DNAJB molecular chaperones to inhibit the aggregation of destabilised client proteins in cells. This Fluc^DM^ isoform acts as a proteostasis sensor by reporting on the capacity of cells to maintain aggregation-prone proteins in a soluble state. Moreover, since Fluc forms amorphous-type aggregates (Schröder et al., 1993; Buchberger et al., 1996; Rampelt et al., 2012) this enabled us to assess whether the mechanism used by DNAJBs to suppress the fibrillar aggregation of proteins (Månsson et al., 2014; Kakkar et al., 2016b) is also used to inhibit the amorphous aggregation of destabilised proteins in cells.

We report that co-expression in cells of DNAJBs with Fluc^DM^ leads to a significant decrease in inclusion formation by Fluc^DM^. Interaction with Hsp70 is required for this activity since mutations that prevent this interaction abolish the capacity of these DNAJBs to suppress the aggregation of Fluc^DM^ into inclusions in cells. Furthermore, our data suggest that DNAJBs support Hsp70-dependent Fluc^DM^ degradation via the proteasome. The S/T-rich domain in DNAJB6 and DNAJB8, which is essential for inhibiting nucleation of polyQ amyloid fibrils (Hageman et al., 2010; Kakkar et al., 2016b), is not required for suppression of Fluc^DM^ aggregation. However, the short TTK-LKS region in the C-terminal of DNAJB8, a region that plays no role in the suppression of polyQ aggregation (Hageman et al., 2010), is essential for anti-aggregation activity against Fluc^DM^. Together, our data suggest that DNAJB6 and DNAJB8 contain at least two domains involved in the inhibition of protein aggregation, an S/T-rich domain for binding β-hairpin structures that go on to form amyloid fibrils and a TTK-LKS region that interacts with exposed areas of hydrophobicity in amorphous substrates. Thus, the intrinsic properties of an aggregation-prone client may dictate the mechanism by which these specialised DNAJBs bind client proteins.

## Materials and methods

### Plasmid constructs

The enhanced green fluorescent protein (EGFP)-N3 plasmid was donated by Dr Darren Saunders (University of New South Wales, Sydney, NSW, Australia). Plasmids encoding wild-type (WT) and double mutant (DM; R188Q, R261Q) Fluc with an N-terminal EGFP tag (Fluc^WT^-EGFP and Fluc^DM^-EGFP) (Gupta et al., 2011) were kindly gifted by Professor Ulrich Hartl (Max Planck Institute of Biochemistry, Munich, Germany) and were cloned into pcDNA4/TO/myc/hisA for mammalian expression by GenScript (Piscataway, NJ, USA). The construction of the V5-tagged DNAJB plasmid library used in this study is described in Hageman and Kampinga (2009). Plasmids expressing pcDNA5/FRT/TO-monomeric red fluorescent protein (mRFP), mutations in DNAJBs in which a histidine residue is replaced with a glutamine (H/Q) within the J-domain and C-terminal deletions in DNAJB8 are outlined in Hageman et al. (2010). Plasmids encoding mutations (M1–4) in the S/T-rich region of DNAJB6 are previously described by Kakkar et al. (2016b) whilst constructs expressing disease-related missense mutations (F93L and P96R) in the G/F-rich region are described by Thiruvalluvan et al. (2020).

### Cell culture, transient transfections and treatment

HEK293 cells (American Type Culture Collection, Manassas, VA, USA) were cultured in DMEM/F-12 supplemented with 2.5 mM L-glutamine (Gibco, Carlsbad, CA, USA) and 10% (v/v) foetal calf serum (FCS) (Gibco) at 37°C under 5% CO2/95% air in a Heracell 150i CO2 incubator (Thermo Fisher Scientific, Glen Burnie, MD, USA). HEK293 stably expressing the tetracycline (tet)-repressor (Flp-In T-REx HEK293, Invitrogen, Carlsbad, CA, USA) were cultured as above with the addition of 50 μg/mL zeocin and 5 μg/mL blasticidin (Invitrogen) to the culture medium weekly to ensure maintenance of the tet-repressor. Cells were routinely tested for mycoplasma contamination (∼ every 6 months) and the identity of these cell lines were verified via short tandem repeat profiling (Garvan Institute of Medical Research, Sydney, NSW, Australia).

For transient transfections, cells were grown to 60-70% confluence in CELLSTAR^®^ 6-well plates (Greiner Bio-One, Frickenhausen, Germany) coated with 0.001% poly-L-lysine (Sigma-Aldrich, St. Louis, MO, USA). Cells were co-transfected 24 h post-plating with linear (MW 25,000) polyethylenimine (BioScientific, Gymea, NSW, Australia) according to the manufacturer’s instructions, and 0.2 µg of plasmid encoding for Fluc^DM^ and 0.8 µg of plasmid DNA encoding for either mRFP (as a negative control) or a DNAJB isoform (WT or mutational variant). For transfection in Flp-In T-REx HEK293 cells, 1 µg/mL tetracycline (Sigma-Aldrich) was added to the culture medium 4 h post-transfection to induce expression. For inhibition of the proteasome, 10 µM MG132 (SelleckChem, Boston, MA, USA) was added to cells 24 h post-transfection and cells were incubated for a further 18 or 24 h. Autophagy was inhibited 24 h post-transfection using a combination of 1 µM bafilomycin A1 (Sapphire Bioscience, Redfern, NSW, Australia) and 5 mM 3-methyladenine (AdipoGen, San Diego, CA, USA) and then the cells were incubated for a further 24 h. Since MG132, bafilomycin A1 and 3-methyladenine were dissolved in dimethyl sulfoxide (DMSO; Sigma-Aldrich), an equivalent volume of DMSO was added to control samples.

### Epifluorescence microscopy

Inclusions formed following expression of Fluc^WT^-EGFP or Fluc^DM^-EGFP were analysed directly in 6-well plates 48 h post-transfection by epifluorescence microscopy. Green fluorescence was detected by excitation at 488 nm. All images were taken at 20X magnification using a Leica DMi8 fluorescence microscope (Leica Microsystems, Wetzlar, Germany). Images were prepared with the Leica Application Suite – Advanced Fluorescence (LAS-AF) Version 3 software (Leica Microsystems).

### Flow cytometry assay to assess inclusion formation

In some experiments, 48 h post-transfection, cells were prepared for flow cytometric analysis using the flow cytometric analysis of inclusions and trafficking (FloIT) method previously described (Whiten et al., 2016). To do so, cells were harvested with 0.05% (v/v) trypsin/EDTA (Gibco), then diluted with DMEM/F-12 containing 1% (v/v) FCS and centrifuged at 300 × *g* for 5 min at RT. Cells were then washed twice in phosphate buffered saline (PBS; 135 mM NaCl, 2.7 mM KCl, 1.75 mM KH2PO4, 10 mM Na2HPO4, pH 7.4) and resuspended in 500 μL PBS. Cells were kept on ice throughout this process to minimise cell death and protein aggregation. An aliquot of the cell suspension (150 μL) was taken and used to measure the transfection efficiency of live cells for use in later analyses.

The remaining cell suspension was centrifuged at 300 × *g* for 5 min at RT and resuspended in 500 μL PBS containing 0.5% (v/v) Triton X-100 (Thermo Fisher Scientific) to facilitate cell lysis. Except in control samples used to set gates, RedDot1 (Biotium, Hayward, CA, USA) was diluted (1:1000) into PBS and then diluted further (1:500) upon addition to cell lysates. Following a 2 min incubation on ice to stain nuclei, flow cytometry was performed as previously described (Whiten et al., 2016). Forward scatter (FSC) and side scatter (SSC), together with RedDot1 fluorescence (640 nm excitation, 670/30 nm collection) and EGFP fluorescence (488 nm excitation, 525/50 nm collection) of particles present in cell lysates were measured. The FSC threshold was set to 200 AU (minimum possible) in order to include small inclusions in the analyses. In all experiments, axes were set to log10 and a minimum of 100,000 events were acquired. Nuclei were identified based on FSC and RedDot1 fluorescence and were excluded from further analyses. Inclusions were counted based on their FSC and EGFP fluorescence, in comparison to untransfected cells. Unless otherwise stated, voltages of 300 (FSC), 200 (SSC), 250, (EGFP) and 550 (RedDot1) were used in all experiments. The number of inclusions identified within the population was normalised against the number of nuclei present and values are reported as the number of inclusions/100 cells according to the equation:

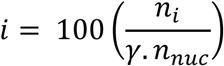

Where n_*i*_ is the number of inclusions present, n_*nuc*_ is the number of nuclei, and *γ* is the transfection efficiency expressed as a fraction (Whiten et al., 2016).

In all experiments, flow cytometry was performed using a BD LSRFortessa X-20 or BD LSR-II analytical flow cytometer (BD Biosciences, San Jose, CA, USA) and FCS files were analysed using FlowJo version 10 (Tree Star Ashland, OR, USA). Histograms were generated and statistical analyses were performed using GraphPad Prism version 8 (GraphPad Software, San Diego, CA, USA). Unless otherwise stated, results are reported as the mean ± standard error of the mean (S.E.M) and the number of independent (biological) replicates (n) of each experiment is specified. Data were analysed by one-way analysis of variance (ANOVA) and Tukey’s post-hoc test or, where appropriate, assessed assuming unequal variance using the two-tailed student t-test. In all analyses, *P* < 0.05 was considered statistically significant.

### Cellular protein extraction, quantification and fractionation

Transfected cells were trypsinised, harvested, washed twice in PBS (300 × *g* for 5 min at RT) and total cellular protein was extracted by lysis with Nonidet™ P-40 (NP-40; Thermo Fisher Scientific) lysis buffer (50 mM Tris-HCl, 150 mM NaCl, 1 mM EDTA, 1% (v/v) NP-40 supplemented with 0.5% (v/v) Halt™ Protease and Phosphatase Inhibitor Cocktail (Thermo Fisher Scientific), pH 8.0). Cell lysates were then sonicated using the Sonifer^®^ 250 Digital cell disruptor and a double step micro-tip (Branson Ultrasonics, Brookfield, CT, USA) at 50% amplitude for 5 sec. The total protein concentration for each sample was then determined using a BCA assay (Thermo Fisher Scientific) according to the manufacturer’s instruction. The concentration in each sample was adjusted with NP-40 lysis buffer to generate cell lysates of 1 mg/mL total protein (total volume was 200 µL) to ensure equal loading onto SDS polyacrylamide gel electrophoresis (SDS-PAGE) gels for subsequent immunoblotting. A 45 µL aliquot of total protein (total fraction) was taken and kept on ice until use. The remaining 155 µL lysate was centrifuged at 20,000 × *g* for 30 mins at 4° C and the supernatant (NP-40 soluble fraction) carefully collected and placed on ice. The pellet was washed in ice-cold TNE buffer (50 mM Tris-HCl, 150 mM NaCl, 1 mM EDTA, pH 8.0) and centrifuged again at 20,000 × *g* for 30 mins at 4° C. The supernatant was carefully removed and discarded and the pellet resuspended in 50 µL NP-40 lysis buffer. The insoluble pellet was sonicated at 50% amplitude for 5 sec (NP-40 insoluble fraction). SDS-PAGE loading buffer (final concentrations: 500 mM Tris-HCl, 2% (w/v) SDS, 25% (w/v) glycerol, 0.01% (w/v) bromophenol blue, 15% (v/v) β-mercaptoethanol (Sigma-Aldrich), pH 6.8) was added to each sample and the samples were then heated at 95°C for 5 min.

### Immunoblotting and detection

Equal amounts of protein were loaded onto 12% (v/v) resolving SDS-PAGE gels with 4% (v/v) polyacrylamide stacking gels using Precision Plus Protein™ dual colour standards (Bio-Rad, Hercules, CA, USA). The gels were run in a Mini-Protean^®^ Tetra Cell system (Bio-Rad) filled with SDS-PAGE running buffer (25 mM Tris base, 192 mM glycine, 0.5% (w/v) SDS, pH 8.3). Samples were electrophoresed for 15 min at 100 V until proteins had migrated through the stacking gel, at which point, the voltage was increased to 150 V and was allowed to run until the bromophenol blue dye front had migrated off the end of the gel (∼1 h). Proteins were transferred onto an ImmunoBlot™ polyvinylidene difluoride (PVDF; Bio-Rad) membrane at 100 V for 1 h in ice-cold transfer buffer (25 mM Tris base, 192 mM glycine, 20% (v/v) methanol, pH 8.3). The membrane was blocked at 4° C overnight with 5% (w/v) skim milk powder in Tris-buffered saline (TBS; 50 mM Tris base, 150 mM NaCl, pH 7.6). Membranes were incubated with either rabbit anti-GFP (1:2500; ab290, Abcam, Cambridge, MA, USA), mouse anti-Hsp70 (1:1000; ab47455, Abcam), mouse anti-SQSTM1/p62 (1:2000; ab56416, Abcam), mouse anti-ubiquitin (1:1000; sc-8017, Santa Cruz Biotechnology, Dallas, TX, USA) or mouse anti-V5 (1:5000; 46-0705, Thermo Fisher Scientific) primary antibodies in 5% (w/v) skim milk powder in TBS-T for 2 h at RT. The membrane was washed four times (each for 10 min) in TBS-T before being incubated with an anti-mouse horse radish peroxidase (HRP)-conjugated secondary antibody (A9044, Sigma-Aldrich) or an anti-rabbit HRP-conjugated secondary antibody (31466, Thermo Fisher Scientific), each diluted 1:5000 into 5% (w/v) skim milk powder in TBS-T. The membrane was rocked at RT for 1 h before being washed four times (each for 10 min) in TBS-T. Proteins of interest were detected with SuperSignal^®^ West Pico Chemiluminescent Substrate or SuperSignal^®^ West Dura Extended Duration Chemiluminescent Substrate (Thermo Fisher Scientific) using an Amersham Imager 600RGB (GE Healthcare Life Sciences, Little Chalfont, UK) or ChemiDoc™ Imaging System (Bio-Rad), with exposure times ranging from 1–15 min.

## Results

### DNAJB family members are potent suppressors of Fluc^DM^ aggregation

We exploited a double mutant isoform of Fluc (Fluc^DM^), which readily forms amorphous aggregates in cells without the need to apply an unfolding stress (Gupta et al., 2011), to determine the capacity of individual DNAJB chaperones to inhibit the aggregation of destabilised proteins. We first confirmed the intracellular aggregation propensity of Fluc^DM^ into inclusions by transfecting cells so they expressed Fluc^DM^-EGFP or the less aggregation-prone Fluc^WT^-EGFP (Fig. 1A). Some cells expressing Fluc^DM^-EGFP contained green fluorescent puncta throughout the cytoplasm (∼15% of cells), corresponding to the aggregation and inclusion formation of this protein. Whilst inclusions were occasionally observed in cells expressing Fluc^WT^-EGFP (less than 5%), most of these cells exhibited diffuse green fluorescence throughout the cytoplasm and the nucleus.

**Figure 1.**
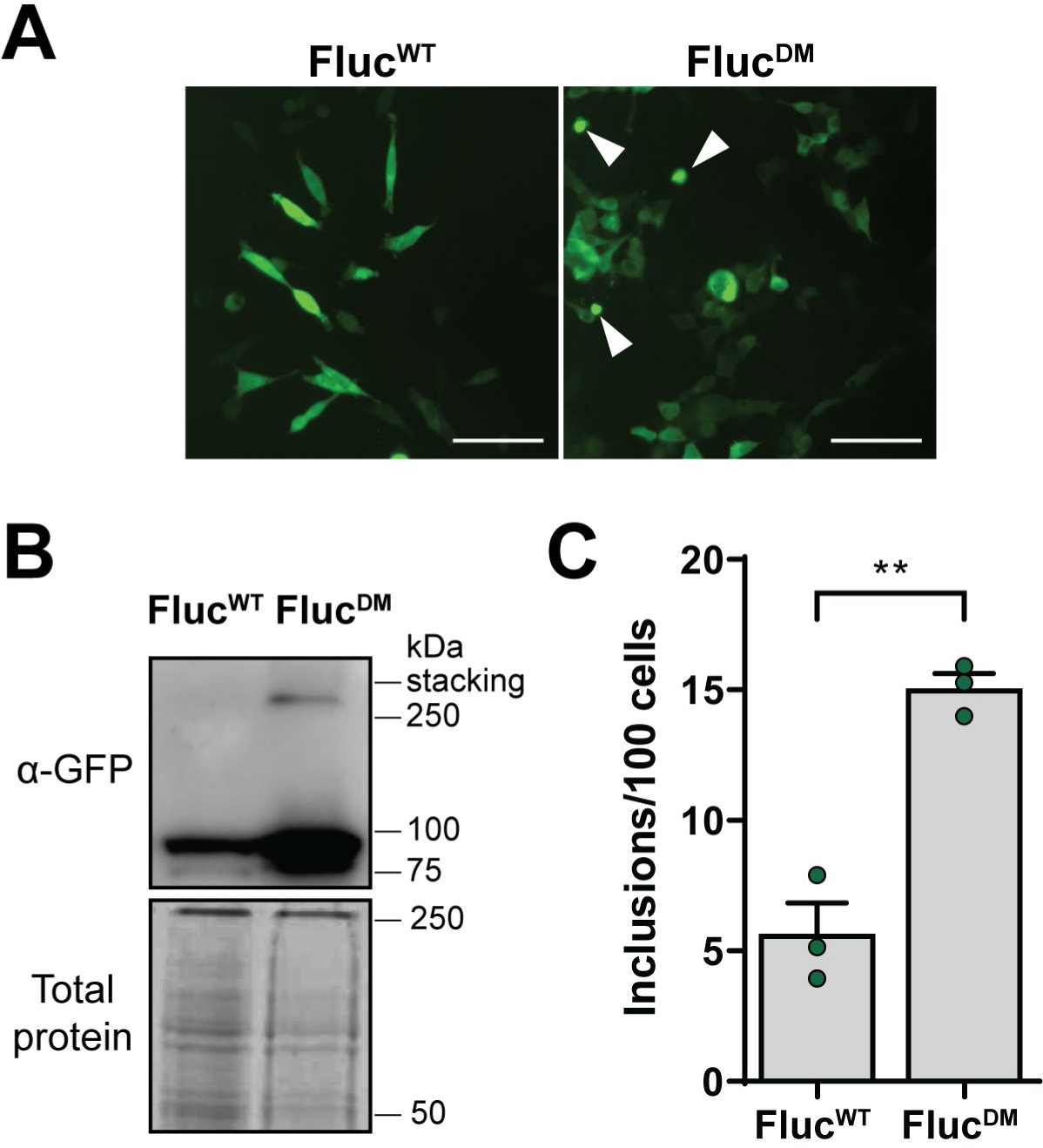
Fluc^DM^ readily aggregates to form inclusions in cells, which can be assessed using FloIT. HEK293 cells were transfected with Fluc^WT^-EGFP or Fluc^DM^-EGFP and analysed 48 h post-transfection by **(A)** epifluorescence microscopy, **(B)** NP-40 cell fractionation followed by immunoblotting or **(C)** quantitative flow cytometry. In (A) green fluorescence was detected by excitation at 488 nm. Examples of cells containing inclusions are denoted by the arrows. All images were taken at 20X magnification using a Leica DMi8 fluorescence microscope. Scale bars = 60 μm. In (B) an anti-GFP antibody was used to detect Fluc^WT^ or Fluc^DM^ in the insoluble pellet fraction and total protein was used as a loading control. Data in (C) is presented as the mean ± S.E.M (n=3) of the number of inclusions per 100 cells. Significant differences between group means in the data were determined using a student’s t-test (** = *P* < 0.01).

The significantly enhanced aggregation propensity of Fluc^DM^-EGFP compared to Fluc^WT^-EGFP was confirmed by examining the distribution of detergent insoluble protein between cells expressing these proteins (Fig. 1B). There was approximately triple the amount of insoluble protein detected in cells expressing Fluc^DM^ compared to Fluc^WT^. Cells were also analysed by the flow cytometric analysis of inclusions and trafficking (FloIT) assay, a technique that readily enumerates the number of inclusions formed in cells (Whiten et al., 2016). The number of inclusions measured by FloIT was significantly higher (∼3 fold) in cells expressing Fluc^DM^ compared to Fluc^WT^ (Fig. 1C), in accordance with the results from the fluorescence microscopy and detergent insolubility of these proteins. Taken together, these data show that Fluc^DM^ readily aggregates in cells and that FloIT can be used as a rapid and non-subjective method to assess this aggregation.

Previous studies have identified that the fibrillar aggregation of polyQ-expanded proteins can be significantly reduced by DNAJB1, DNAJB6b and DNAJB8, while other DNAJBs are much less effective (Hageman et al., 2010). To test the ability of DNAJBs to engage destabilised proteins at risk of forming amorphous aggregates in cells, V5-tagged DNAJBs were transiently co-expressed with Fluc^DM^ in Flp-In T-REx HEK293 cells. Using the FloIT assay to assess the aggregation of Fluc^DM^, we showed that inclusion formation was significantly reduced by all DNAJBs tested, with DNAJB1, DNAJB5, DNAJB6b and DNAJB8 being the most potent suppressors of inclusion formation (Fig. 2).

**Figure 2.**
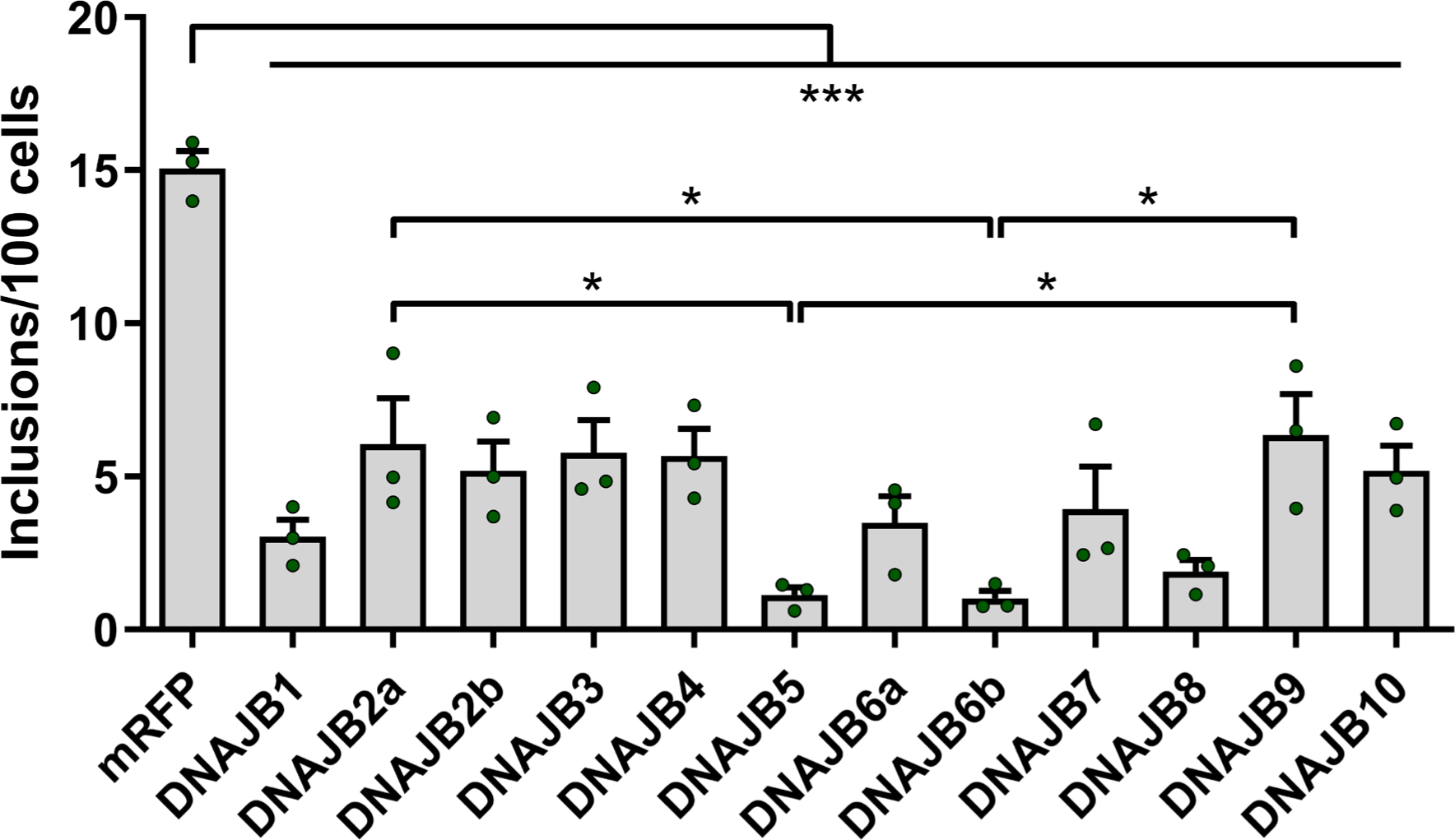
DNAJBs prevent the intracellular aggregation of Fluc^DM^ into inclusions. Flp-In T-REx HEK293 cells were co-transfected with V5-tagged DNAJBs (or mRFP as a negative control) and Fluc^DM^-EGFP. Expression of DNAJBs was induced by addition of tetracycline and whole cell lysates were analysed by FloIT 48 h post-transfection. Data is presented as the mean ± S.E.M (n=3) of the number of inclusions per 100 cells. Significant differences between group means in the data were determined using a one-way ANOVA (*P* < 0.05) followed by a Tukey’s post-hoc test. Group means determined to be statistically different from each other are indicated (\**P* < 0.05 and \*\*\**P* < 0.001).

Our findings that all of the DNAJBs tested significantly suppressed Fluc^DM^ aggregation contrasts to the ability of only a few specific DNAJB isoforms (DNAJB2a, DNAJB6 and DNAJB8) to strongly inhibit polyQ aggregation, with DNAJB1 having an intermediate effect (Hageman et al., 2010). However, unlike what we have observed for suppression of inclusion formation by Fluc^DM^, DNAJB4, DNAJB5, DNAJB9 were significantly less active against polyQ aggregation, with DNAJB2b having no effect at all (Hageman et al., 2010). This suggests that the mechanism by which DNAJBs act to suppress the amorphous aggregation of Fluc^DM^ is not the same as that used to suppress the fibrillar aggregation of proteins. Since DNAJB1 (weak polyQ aggregation inhibitor) and DNAJB6b (herein referred to as DNAJB6) and DNAJB8 (strong polyQ aggregation inhibitors) were among the most effective DNAJBs at suppressing Fluc^DM^ aggregation, we focussed on these isoforms in order to further interrogate this mechanism of action.

### DNAJBs promote the degradation of Fluc^DM^, primarily via the proteasome

We sought to determine whether the inhibition of Fluc^DM^ aggregation into inclusions by DNAJBs requires the degradative activity of the proteasome or autophagy. To do so, HEK293 cells were co-transfected to express Fluc^DM^ and DNAJB1 or DNAJB6 (or mRFP as a control) and, 24 h post-transfection, were treated with proteasome (MG132) or autophagy (3-methyladenine + bafilomycin A1) inhibitors and analysed at 48 h. Treatment of cells with inhibitors of autophagy had no significant effect on the level of inclusion formation and little effect on the insolubility of Fluc^DM^ in cells co-expressing a DNAJB (Fig. 3A,B). Inhibition of autophagy was confirmed in cells treated with 3-methyladenine + bafilomycin A1 by the increased levels of SQSTM1/p62 (a commonly used marker of autophagy). An increase in SQSTM1/p62 was also observed in cells treated with MG132, an effect which has been reported previously in the literature whereby inhibition of the proteasome leads to upregulation of p62 transcription (Myeku and Figueiredo-Pereira, 2011), suggesting a crosstalk between the two pathways (Liu et al., 2016). As there was no substantial increase in the level of insoluble Fluc^DM^ in cells treated with the autophagy inhibitors, these data suggest that Fluc^DM^ is primarily degraded by the proteasome.

**Figure 3.**
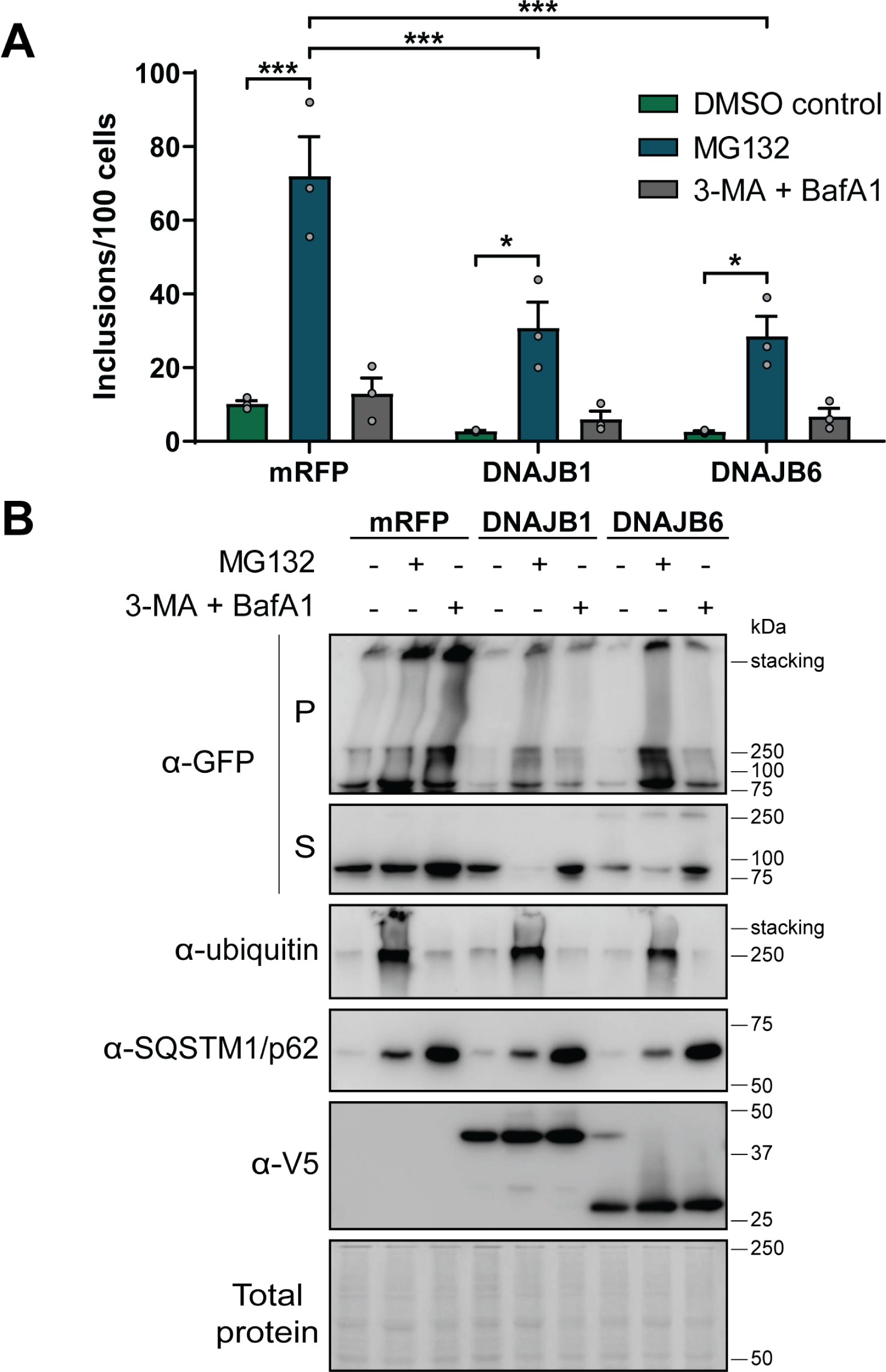
DNAJBs require an active proteasome to facilitate the degradation of Fluc^DM^. HEK293 cells were co-transfected to express Fluc^DM^-EGFP and mRFP (as a negative control), DNAJB1 or DNAJB6 and 24 h post-transfection, cells were treated with the proteasome inhibitor MG132 (10µM) or autophagy inhibitors 3-methyladenine (5mM) and bafilomycin A1 (1µM), or a DMSO vehicle control. Cells were incubated for a further 24 h and then analysed by **(A)** quantitative flow cytometry or **(B)** NP-40 fractionation and subsequent immunoblotting. Data in (A) is presented as the mean ± S.E.M (n=3) of the number of inclusions per 100 cells. Significant differences between group means in the data were determined using a one-way ANOVA (*P* < 0.05) followed by a Tukey’s post-hoc test. Group means determined to be statistically different from each other are indicated (\**P* < 0.05 and \*\*\**P* < 0.001). In (B) an anti-GFP antibody was used to detect Fluc^DM^ in the insoluble pellet (P) and soluble (S) fractions. In the total protein fraction, the expression of DNAJBs were detected with an anti-V5 antibody, an anti-ubiquitin antibody was used to detect ubiquitinated proteins and an anti-SQSTM1/p62 antibody was used to assess autophagy inhibition. Total protein was used as a loading control.

Upon treatment with MG132, the number of Fluc^DM^ inclusions as assessed by FloIT significantly increased in cells co-expressing the mRFP non-chaperone control (Fig. 3A) and this corresponded to an increase in the proportion of Fluc^DM^ found in the NP-40 insoluble fraction (Fig. 3B). Inhibition of the proteasome following treatment with MG132 was evidenced by large smears of polyubiquitinated protein in these samples. Proteasome inhibition led to a significant increase in inclusion formation in cells overexpressing DNAJBs compared to vehicle (DMSO) treated cells, such that the capacity of co-expressed DNAJBs to reduce the amount of insoluble Fluc^DM^ formed was significantly reduced when cells were treated with MG132. Additionally, the amount of soluble Fluc^DM^ also decreased in cells expressing DNAJBs treated with MG132 compared to DMSO controls. This was despite aggregation still being significantly reduced in the DNAJB overexpressing cells under these proteasomal impaired conditions when compared to cells not expressing additional DNAJBs. The DNAJBs can keep the misfolded protein in a non-aggregated soluble form (proteasome independent) such that, with time, the proteasome can facilitate their degradation. Thus, together these data indicate that Fluc^DM^ aggregation is dependent upon proteasomal degradation and that the inhibition of Fluc^DM^ aggregation into inclusions by DNAJBs requires the activity of the proteasome.

### The J-domain is crucial for DNAJBs to protect against Fluc^DM^ aggregation

We next examined whether DNAJBs require an interaction with Hsp70 in order to suppress the aggregation of Fluc^DM^ into inclusions. To do so, we employed mutant forms of the DNAJBs in which a histidine residue is replaced with a glutamine (H/Q) within the highly conserved histidine-proline-aspartate (HPD) motif of the J-domain (Hageman et al., 2010) (Fig. 4A). The HPD motif plays a critical role in the regulation of Hsp70 activity; the H/Q mutation in this motif blocks the ability of the DNAJB to interact with Hsp70 (Cheetham and Caplan, 1998), thereby abrogating its ability to stimulate Hsp70 ATPase activity (Tsai and Douglas, 1996) and recruit Hsp70 to clients. The H/Q mutation abolished the capacity of each of the three DNAJBs to inhibit the aggregation of Fluc^DM^, as evidenced by FloIT and assessment of the aggregation of Fluc^DM^ by NP-40 cell fractionation (Fig. 4B,C). Thus, co-expression of the WT DNAJBs reduced the amount of insoluble protein whereas cells expressing the H/Q mutant isoforms contained an equivalent or increased amount of insoluble Fluc^DM^ compared to the mRFP control. The amount of soluble Fluc^DM^ in cells expressing a WT DNAJB decreased compared to cells expressing the mRFP control. This effect is likely due to there being less total Fluc^DM^ in cells expressing WT DNAJBs due to them promoting its degradation. The expression of the H/Q variants were slightly higher than the corresponding WT protein and this could be due to the mutant becoming trapped with their substrates within inclusions, such that their own normal turnover is delayed. Strikingly, the relative loss in activity of the H/Q variants was highest for DNAJB1 (i.e. largest increase in insoluble protein compared to WT variant) and the distribution of insoluble to soluble Fluc^DM^ was different to that of cells expressing DNAJB6 H/Q or DNAJB8 H/Q. Given the expression of endogenous Hsp70 was unaffected under these experimental conditions, these data imply that all DNAJBs prevent the aggregation of Fluc^DM^ by interacting with Hsp70 but that their mode of action and their Hsp70 dependence may be dissimilar.

**Figure 4.**
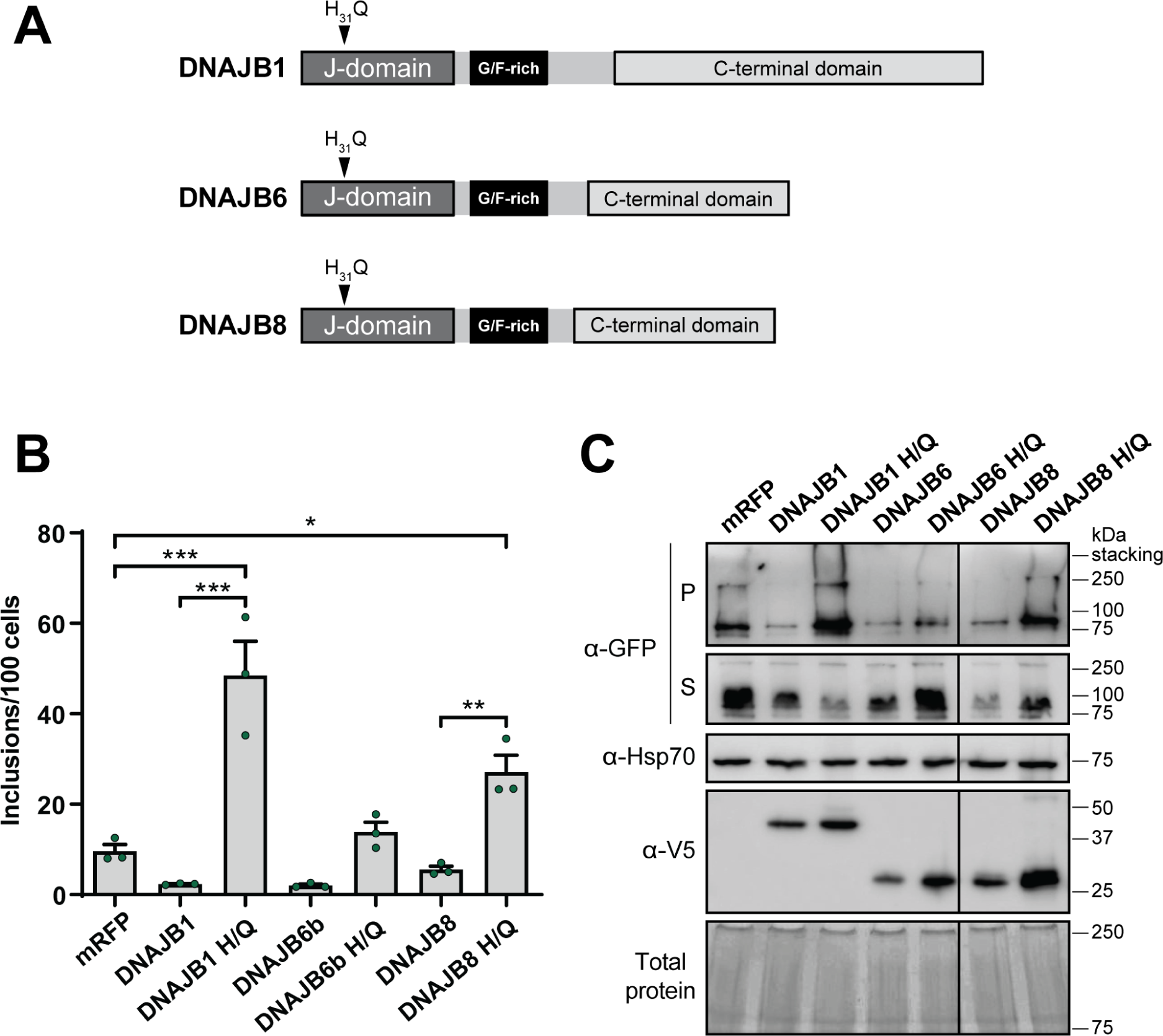
Interaction with Hsp70 is required for DNAJBs to suppress Fluc^DM^ aggregation. **(A)** Schematic overview of DNAJB proteins identifying location of mutation within the J-domain, in which the histidine residue has been substituted for a glutamine (termed H/Q) at amino acid position 31 within the HPD (Hsp70-interacting) motif. HEK293 cells were co-transfected to express Fluc^DM^-EGFP and mRFP (as a negative control), DNAJB1, DNAJB6, DNAJB8 or their H/Q variants. Cells were analysed 48 h post-transfection by **(B)** quantitative flow cytometry or **(C)** NP-40 cell fractionation followed by immunoblotting. Data in (B) is presented as the mean ± S.E.M (n=3) of the number of inclusions per 100 cells. Significant differences between group means in the data were determined using a one-way ANOVA *(P* < 0.05) followed by a Tukey’s post-hoc test. Group means determined to be statistically different from each other are indicated (\**P* < 0.05, \*\**P* < 0.01 and \*\*\**P* < 0.001). In (C) an anti-GFP antibody was used to detect Fluc^DM^ in the insoluble pellet (P) and soluble (S) fractions. In the total protein fraction, the expression of DNAJBs was detected with an anti-V5 antibody and an anti-Hsp70 was used to detect endogenous expression of Hsp70. Total protein was used as a loading control.

### DNAJBs facilitate interaction with Hsp70 and Fluc^DM^ for proteasomal degradation

In order to examine whether DNAJBs mediate Fluc^DM^ degradation by the proteasome via interaction with Hsp70, we co-expressed Fluc^DM^ and the DNAJB H/Q variants in cells and then treated with MG132. We surmised that if Hsp70 was the driver of proteasomal degradation of Fluc^DM^, the H/Q mutants, which are unable to interact with Hsp70, should not further increase the levels of inclusions formed in MG132-treated cells. Inhibition of the proteasome in these experiments was again confirmed by an increase in polyubiquitinated species, observed as large, high molecular weight smears by immunoblotting with an anti-ubiquitin antibody. As before, there was a significant increase in the number of inclusions in cells expressing either the DNAJB1 H/Q or DNAJB8 H/Q variant compared to the mRFP expressing control (Fig. 5A). However, inclusion formation did not further increase in cells expressing an H/Q variant which was treated with MG132. Again, the result was different for DNAJB1 compared to DNAJB6 and DNAJB8, whereby treatment of cells expressing DNAJB1with MG132 lead to a decline in the number of inclusions compared to the DMSO vehicle control. There was no difference in the amount of insoluble protein detected between cells expressing the H/Q variants treated with MG132 compared to vehicle-treated (DMSO) controls (Fig. 5B). We did note some inter-assay variability for cells expressing mRFP treated with MG132 compared to previous experiments (Fig. 3A); we attribute this to differences in the time cells were treated with MG132 (i.e. cells were treated with MG132 for 24 h in the experiments presented in Fig. 3A and 18 h in the experiments presented in Fig. 5A). Taken together, these data provide further evidence that all DNAJBs antagonise Fluc^DM^ aggregation by keeping it competent for proteasomal degradation, which requires interaction with Hsp70 to be effective.

**Figure 5.**
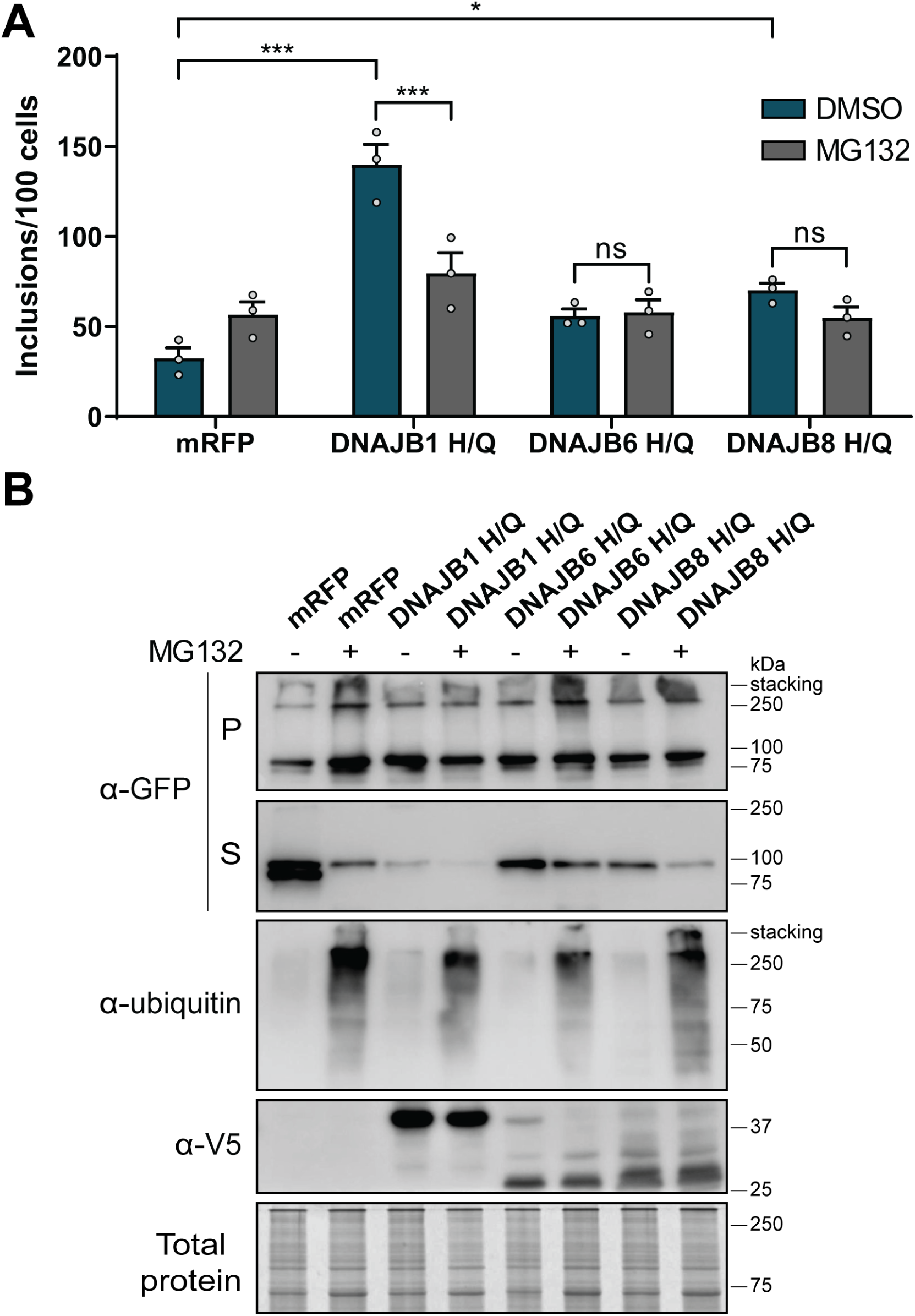
DNAJBs rely upon interaction with Hsp70 to deliver Fluc^DM^ for the degradation via the proteasome. HEK293 cells co-transfected with Fluc^DM^-EGFP and mRFP (as a negative control), DNAJB1, DNAJB6 or DNAJB8 H/Q variants. Cells were treated with a proteasome inhibitor MG132 (10µM) or a DMSO vehicle control 24 h post-transfection. Cells were incubated for a further 18 h and analysed 42 h post-transfection by **(A)** quantitative flow cytometry or **(B)** NP-40 fractionation and subsequent Western blotting. Data in (A) is presented as the mean ± S.E.M (n=3) of the number of inclusions per 100 cells. Significant differences between group means in the data were determined using a one-way ANOVA (*P* < 0.05) followed by a Tukey’s post-hoc test. Group means which are statistically significant are indicated (ns represent non-significant groups, \**P* < 0.05 and \*\*\**P* < 0.001). In (B) an anti-GFP antibody was used to detect Fluc^DM^ in the insoluble pellet (P) and soluble (S) fractions. In the total protein fraction, expression of DNAJBs was detected with an anti-V5 antibody and an anti-ubiquitin antibody was used to observe inhibition of the proteasome. Total protein was used as a loading control.

### Disease-related mutations in the G/F-rich region of DNAJB6 do not impact the capacity to prevent the aggregation of Fluc^DM^ into inclusions

To probe for other regions within DNAJBs that are required for suppressing Fluc^DM^ aggregation, we first assessed the impact of two disease-related missense mutations within the G/F-rich region of DNAJB6 (F93L and P96R) (Fig. 6A). The F93L and P96R mutations have been associated with limb-girdle muscular dystrophy and it has been suggested that these mutations lead to disruption of the J to G/F-interdomain interaction and minor loss of function in their capacity to suppress polyQ aggregation (Sarparanta et al., 2012; Thiruvalluvan et al., 2020). However, we found that both the F93L and P96R mutational variants of DNAJB6 fully retained the ability to inhibit the aggregation of destabilised Fluc^DM^ into inclusions (Fig. 6B,C).

**Figure 6.**
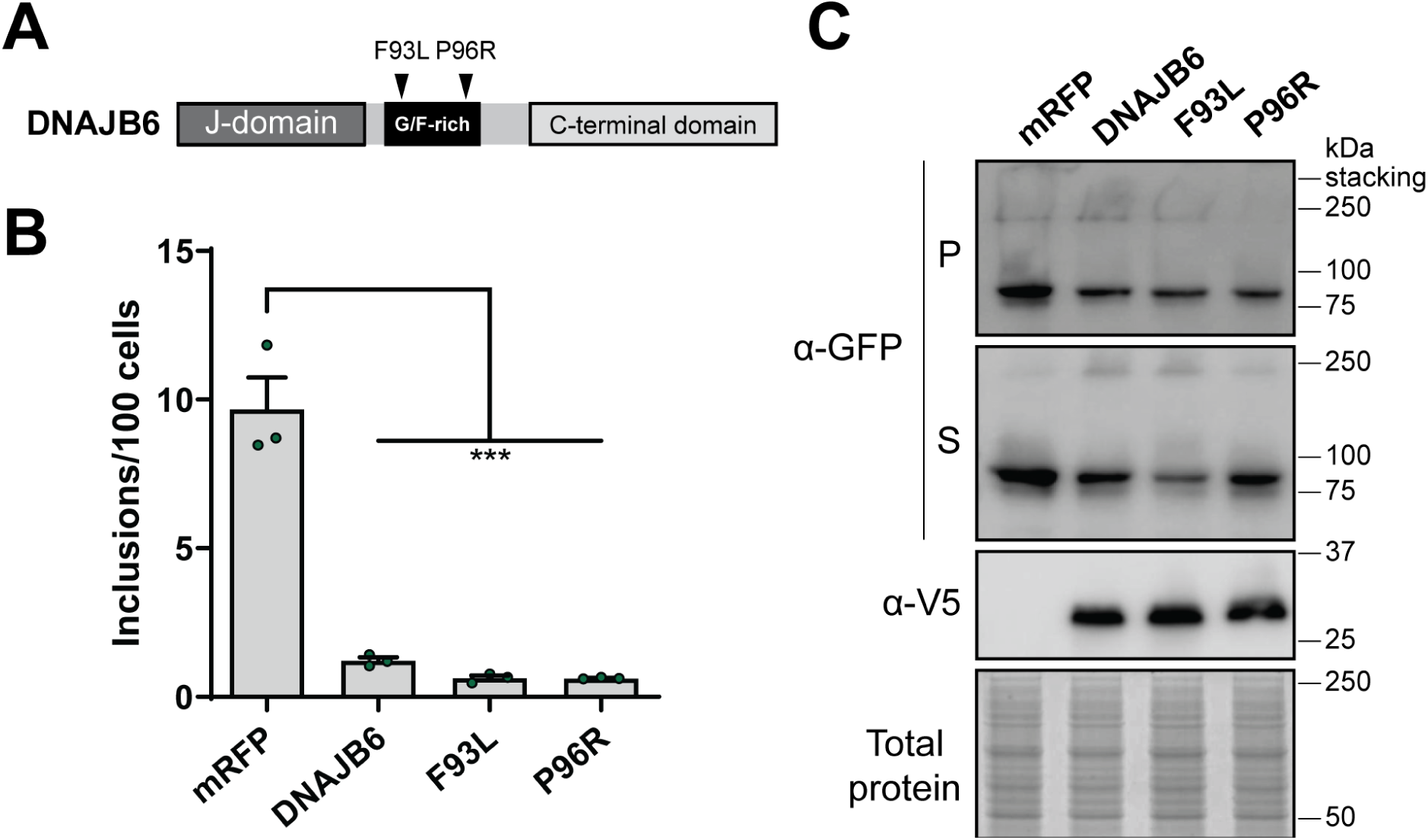
Disease-related mutations in the G/F-rich domain of DNAJB6 do not affect its capacity to inhibit Fluc^DM^ inclusions formation. **(A)** Schematic overview of DNAJB6 disease-related missense mutations at amino acid positions 93 and 96 in the G/F-rich region. HEK293 cells were co-transfected with Fluc^DM^-EGFP and mRFP (as a negative control) or DNAJB6 G/F-domain disease-related mutational variants. Cells were analysed 48 h post-transfection by **(B)** quantitative flow cytometry or **(C)** NP-40 fractionation and immunoblotting. Data in (B) is presented as the mean ± S.E.M (n=3) of the number of inclusions per 100 cells. Significant differences between group means in the data were determined using a one-way ANOVA (*P* < 0.05) followed by a Tukey’s post-hoc test. Group means determined to be statistically different from each other are indicated (\*\*\**P* < 0.001). In (C) an anti-GFP antibody was used to detect Fluc^DM^ in the insoluble pellet (P) and soluble (S) fractions. The expression of DNAJBs was detected with an anti-V5 antibody in the total protein fraction. Total protein was used as a loading control.

### The C-terminus, and not the serine-rich region of DNAJBs, is required for DNAJBs to suppress Fluc^DM^ inclusion formation in cells

Previous work has suggested that the hydroxyl groups of serine/threonine (S/T) side chains in the C-terminal domain of DNAJB6 participate in intramolecular hydrogen bonding with polyQ peptides and that this likely mediates inhibition of amyloid formation. For example, increasing the number of S/T residues substituted with alanine (A) residues (from 6, to 13 and to 18 substitutions; variants referred to as M1, M2 or M3, respectively: Fig. 7A), leads to a progressive loss in the ability of DNAJB6 to inhibit polyQ or amyloid-β aggregation, with the M3 variant being rendered completely functionally inactive (Kakkar et al., 2016b; Månsson et al., 2018). Interestingly, we found that the DNAJB6 M1, M2 and M3 variants fully retain their ability to suppress intracellular inclusion formation by Fluc^DM^ (Fig. 7B and C), indicating that the hydroxyl groups of the S/T-rich domain of DNAJB6 are not required for interaction between DNAJB6 and Fluc^DM^. Deletion of almost the entire S/T-rich region of the C-terminus of DNAJB6 (M4) did result in abrogation of DNAJB6-mediated suppression of Fluc^DM^ aggregation; however, this is likely due to structural destabilisation of this DNAJB6 mutant which results in it being readily degraded (Kakkar et al., 2016b), as evidenced by its very low levels in the lysate from transfected cells. Together, these data imply that the residues in DNAJB6 responsible for the inhibition of the amorphous aggregation of Fluc^DM^ into inclusions differ from those used to suppress amyloid fibril-type aggregation of proteins.

**Figure 7.**
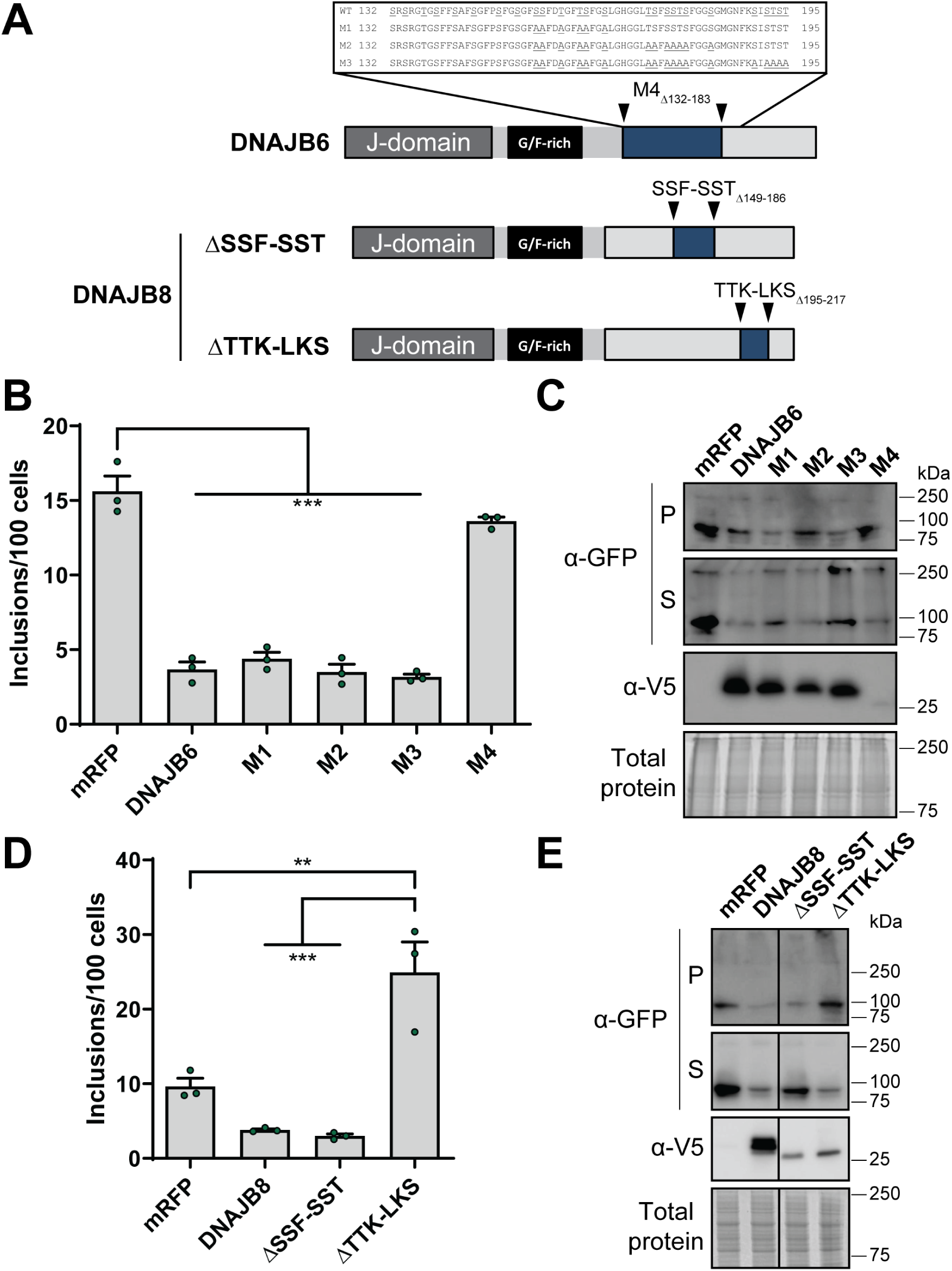
The TTK-LKS region of C-terminus of DNAJB8 is required to suppress the aggregation of Fluc^DM^into inclusions. **(A)** Schematic overview of DNAJB6 and DNAJB8 C-terminal mutational variants used in this work. M1, M2 and M3 are mutations in the S/T-rich region of DNAJB6 in which underlined amino acids represent 6, 13 and 18 S/T-to-A substitutions, respectively. Regions identified between sets of arrows indicate deletion mutations. HEK293 cells were co-transfected with Fluc^DM^-EGFP and DNAJB6 or DNAJB8 C-terminal mutational variants (or mRFP as a negative control) and analysed 48 h post-transfection by quantitative flow cytometry **(B and D)** or NP-40 fractionation and subsequent immunoblotting **(C and E)**. Data in (B and D) is presented as the mean ± S.E.M (n=3) of the number of inclusions per 100 cells. Significant differences between group means in the data were determined using a one-way ANOVA *(P* < 0.05) followed by a Tukey’s post-hoc test. Group means determined to be statistically different from each other are indicated (\*\**P* < 0.01 and \*\*\**P* < 0.001). In (C and E) an anti-GFP antibody was used to detect Fluc^DM^ in the insoluble pellet (P) and soluble (S) fractions. The expression of DNAJBs was detected with an anti-V5 antibody in the total protein fraction. Total protein was used as a loading control.

To further define the functional regions within the C-terminal domain responsible for the anti-aggregation activity of DNAJBs, two DNAJB8 deletion constructs were co-expressed with Fluc^DM^ in cells (Fig. 7A). In accordance with the results obtained by expression of the M3 isoform of DNAJB6, following deletion of the S/T-rich region (ΔSSF-SST), DNAJB8 retained the ability to suppress the aggregation of Fluc^DM^, as determined by both quantitative flow cytometry and cell fractionation (Fig. 7D,E). This again is in contrast to what has been reported previously with regards to the regions in DNAJB8 required for inhibiting the fibrillar aggregation of polyQ (Hageman et al., 2010). Conversely, deletion of the short C-terminal TTK-LKS motif, which is dispensable for inhibiting polyQ aggregation (Hageman et al., 2010), completely abrogates the capacity of DNAJB8 to inhibit Fluc^DM^ inclusion formation (Fig. 7D,E). Together these data suggest that DNAJB6 and DNAJB8 may have at least two different regions involved in substrate handling, one that is responsible for proteins that form β-hairpins during amyloid formation (Kakkar et al., 2016b) and another that is required for the handling of destabilised aggregation-prone proteins, such as those represented here by Fluc^DM^.

## Discussion

In this work, we demonstrate that the DNAJB molecular chaperones are potent suppressors of the aggregation of Fluc^DM^ into inclusions in cells. This contrasts with what has been observed previously, whereby only specific DNAJB isoforms suppress the aggregation of polyQ-expanded proteins (Hageman et al., 2010). For DNAJB1, DNAJB6 and DNAJB8, we show that they inhibit Fluc^DM^ aggregation in a manner that depends on their ability to interact with Hsp70 and is associated with the cellular capacity to degrade Fluc^DM^ via the proteasome, thereby alleviating protein aggregation. For DNAJB6 and DNAJB8, the suppression of Fluc^DM^ aggregation not only appears mechanistically different from DNAJB1, but is also distinct from what has been reported previously for the handling of amyloid-fibril forming proteins, such as polyQ-expanded proteins and the amyloid-β peptide (Hageman et al., 2010; Kakkar et al., 2016b; Månsson et al., 2018). Whilst the S/T-rich region and, to some extent, the G/F-rich region (Sarparanta et al., 2012; Thiruvalluvan et al., 2020) in DNAJB6 and DNAJB8 are essential for amyloid suppression, these regions are not required to inhibit Fluc^DM^ inclusion formation. Finally, we identified a short, 23 amino acid (TTK-LKS) sequence in the C-terminus of DNAJB8 that is required to inhibit Fluc^DM^ inclusion formation even though it is was found to be dispensable for suppression of polyQ aggregation (Hageman et al., 2010). Thus, whilst DNAJB6 and DNAJB8 are both potent inhibitors of polyQ and Fluc^DM^ aggregation, the mechanism by which they interact with these aggregation-prone proteins is different. Our data suggests that DNAJB6 and DNAJB8-like proteins possess distinct regions for binding clients and that this is likely dictated by the structure or composition of the aggregation-prone protein.

We exploited the previously described flow cytometry technique FloIT (Whiten et al., 2016) to screen the capacity of various DNAJB isoforms to suppress Fluc^DM^ inclusion formation. Previous to this, Hsp overexpression screens to identify inhibitors of protein aggregation have typically relied on using traditional bulk-based biochemical analyses, such as the filter trap assay or fluorescence microscopy (Hageman et al., 2010; Kakkar et al., 2016a; Serlidaki et al., 2020). However, when screening many different samples, these types of assays can be time consuming, laborious and, when it comes to counting inclusions in individual cells, subjective. Whilst the basic principle behind methods such as the filter trap assay and FloIT are the same (i.e. analysis of insoluble protein in a cell lysate), FloIT offers several advantages. First, FloIT can be used in a medium to high throughput capacity for rapid non-subjective quantification of inclusions across samples that may differ in transfection efficiency and cell number. Second, FloIT can identify (and enumerate) inclusions that differ in size, granularity and even protein composition, making FloIT broadly applicable to most (if not all) model systems of protein aggregation in cells (Whiten et al., 2016). Here, we describe the first use of FloIT as a quantitative method to screen for the ability of molecular chaperones (and mutational variants) to prevent protein aggregation in cells, highlighting the power and potential applications of the technique to the proteostasis field.

The S/T-rich stretch in DNAJB6 (amino acids 155-195) and DNAJB8 (amino acids 149-186) is highly conserved between these proteins. It has been proposed that interaction with hydroxyl groups in side chains of these S/T residues inhibits primary nucleation by outcompeting for hydrogen bonding essential for β-hairpin and mature amyloid fibril formation, thereby suppressing aggregation (Kakkar et al., 2016b). DNAJB2 also contains a partial serine-rich stretch and, although it is not confirmed to be involved in polyQ handling, is also more effective than DNAJB1 (which lacks this region) at suppressing polyQ aggregation (Hageman et al., 2010). However, mutation or deletion of this S/T-rich region did not result in loss of the ability of DNAJB6 or DNAJB8 to suppress the aggregation of Fluc^DM^ into inclusions, indicating that a different region of the protein is involved in this process. Interestingly, DNAJB1 and the other DNAJBs we tested were all capable of suppressing Fluc^DM^ aggregation. Whilst our data cannot provide insight into the domains required by these other DNAJBs to prevent Fluc^DM^ aggregation, our work has identified a short TTK-LKS (TTKRIVENGQERVEVEEDGQLKS) fragment conserved between DNAJB6 (amino acids 204-226) and DNAJB8 (amino acids 195-217) that is crucial for handling of Fluc^DM^. Since this TTK-LKS domain in DNAJB8 is dispensable for its the capacity to inhibit polyQ aggregation (Hageman et al., 2010), our findings are the first to demonstrate that DNAJB8 has two distinct client interaction domains. Little is currently known regarding the functional role of this conserved TTK-LKS motif in DNAJB6 and DNAJB8. However, based on our current data, we hypothesise that it is either directly or indirectly involved in binding hydrophobic patches in destabilised aggregation-prone proteins.

Recent structural homology modelling of the DNAJB6 dimer/oligomer revealed four β-strands within the C-terminal domain of DNAJB6 (Söderberg et al., 2018). Dimerisation of each DNAJB6 monomer likely occurs via same-to-same-residue crosslinks at lysine residues K189 and K232 within the first and fourth β-strands, respectively. When cross-linked to form a dimer, the symmetrically positioned β-strands within DNAJB6 monomers form a peptide-binding pocket which is surface exposed and lined with the S/T residues responsible for binding fibrillar proteins. Based on these structural data, the TTK-LKS region (which lies downstream of the S/T-rich region) is contained within the fourth β-strand of DNAJB6/8, which is surface-exposed in the monomeric and dimeric form of DNAJB6. Thus, this region has the potential to be a second substrate-binding region in DNAJB6/8, responsible for binding hydrophobic aggregation-prone client proteins

One possible reason why DNAJB6 and DNAJB8-like proteins possess distinct mechanisms to interact with aggregation-prone client proteins is due to intrinsic structural differences in misfolded states of proteins that lead to the formation of amorphous aggregates as opposed to amyloid fibrils. PolyQ-expanded proteins form large, tightly aggregated structures that are extremely insoluble and typical of amyloidogenic deposits (Hageman et al., 2010; Kubota et al., 2011). The R188Q and R261Q mutations in the N-terminus of the mutant Fluc model protein used in this study conformationally destabilises the protein (Gupta et al., 2011), thereby inducing protein misfolding and increased regions of exposed hydrophobicity. This causes the protein to form aggregates that are SDS-soluble and localise into diffuse cytosolic inclusions (Gupta et al., 2011), distinct from the amyloid-like aggregates formed by polyQ-expanded proteins. Indeed, when both polyQ-expanded huntingtin and Fluc^DM^ are expressed together in human cell lines, the two proteins deposit into distinct aggregated structures (Gupta et al., 2011), reaffirming that they aggregate via different mechanisms. Importantly, our data highlight that it may be possible to design therapeutics that boost the ability of DNAJB6/8 to prevent amyloid fibril formation associated with disease, whilst not impacting its capacity to interact with highly destabilised aggregation-prone proteins destined for degradation by the proteasome.

In conclusion, we have utilised the proteostasis sensor Fluc^DM^ to show that overexpression of the DNAJB molecular chaperones acts to boost the PQC capacity of cells. We demonstrate that the ability of DNAJBs to inhibit the aggregation of Fluc^DM^ into inclusions relies on interaction with Hsp70 and this facilitates degradation of Fluc^DM^ by the proteasome. Significantly, we show that the TTK-LKS region in the C-terminal domain of DNAJB6/8 is essential for engaging this destabilised client protein and preventing its aggregation. Moreover, we show that the S/T-rich region of DNAJB6-like proteins that mediates interactions with amyloid forming client proteins is not involved in the suppressing Fluc^DM^ aggregation. Our data highlights the important role of DNAJB molecular chaperones in preventing all forms of protein aggregation in cells and highlights the potential of targeting them for the amelioration of diseases associated with protein aggregation.

## Acknowledgements

We thank Ms Maria van Waarde-Verhagen for technical assistance and staff in Molecular Horizons and the Illawarra Heath and Medical Research Institute for technical and administrative support.

## List of abbreviations

**Abbreviation Definition**

ALS: Amyotrophic lateral sclerosis
ANOVA: Analysis of variance
EGFP: Enhanced green fluorescent protein
FCS: Foetal calf serum
FloIT: Flow cytometric analysis of inclusions and trafficking
Fluc: Firefly luciferase
FlucDM: Firefly luciferase double mutant
FlucWT: Firefly luciferase wild-type
Hsp: Heat shock protein
HspmRFP: Monomeric red fluorescent protein
HspPBS: Phosphate buffered saline
HspPolyQ: Polyglutamine
HspPQC: Protein quality control
HspProteostasis: Protein homeostasis
HspSOD1: Superoxide dismutase 1
HspTDP-43: TAR DNA-binding protein 43

## Conflict of interest

The authors declare no conflict of interest.

## Author contributions

S.M., H.E., S.B. and H.H.K conceptualised the project and the experimental approach. S.M. performed the experiments, analysed the data, constructed the figures and wrote the initial manuscript. H.E., S.B. and H.H.K edited the manuscript and approved the submission of the final manuscript.

## Funding

S.M. was supported with an Australian Government Research Training Program Scholarship and a New Holland Scholarship presented by Nuffic. This work was supported by grants from Science without Borders from the Brazilian Government (to H.H.K.), the Dutch Campaign Team Huntington (to S.B. and H.H.K.) and The Netherlands Organization for Scientific Research (ZonMw; project numbers 733051076 and 91217002 to H.H.K).

## Notes

### Competing Interest Statement

The authors have declared no competing interest.

